# G9a/GLP-Sensitivity of H3K9me2 Demarcates Two Types of Genomic Compartments

**DOI:** 10.1101/2020.06.26.173849

**Authors:** Zixiang Yan, Luzhang Ji, Xiangru Huo, Qianfeng Wang, Yuwen Zhang, Bo Wen

## Abstract

In the nucleus, chromatin is folded into hierarchical architecture that is tightly linked to various nuclear functions. However, the underlying molecular mechanisms that confer these architectures remain incompletely understood. Here, we investigated the functional roles of H3 lysine 9 dimethylation (H3K9me2), one of the abundant histone modifications, in three-dimensional (3D) genome organization. Unlike mouse embryonic stem cells (mESCs), inhibition of methyltransferases G9a and GLP in differentiated cells eliminated H3K9me2 predominantly at A-type (active) genomic compartments, and the level of residual H3K9me2 modification was strongly associated with genomic compartments in differentiated cells. Furthermore, chemical inhibition of G9a/GLP in mouse hepatocytes led to the decreased chromatin-nuclear lamina interactions mainly at G9a/GLP sensitive regions (GSRs), the increased degree of genomic compartmentalization, and the up-regulation of hundreds of genes that were associated with alterations of the 3D chromatin. Collectively, our data demonstrated essential roles of H3K9me2 in 3D genome organization.

## Introduction

The H3 lysine 9 dimethylation (H3K9me2) is one of the histone modifications involved in gene silencing and repressive chromatin [1]. Unlike H3K9me3 that associates with constitutively pericentromeric heterochromatin, H3K9me2 marks broad chromatin domains that are named large organized chromatin K9 modifications (LOCKs) across the genome [2, 3]. H3K9me2 is mainly deposited by methyltransferases G9a (EHMT2) and GLP (EHMT1), which primarily form stable heterodimer *in vivo*. G9a and GLP cannot compensate each other, and depletion of either can significantly reduce the level of H3K9me2; a double knockout, however, does not get further effect [4, 5]. H3K9 methylation is selectively recognized and bound by heterochromatin protein 1 (HP1) family members, HP1α, HP1β, and HP1γ (also known as CBX5, CBX1, and CBX3, respectively), which functions to promote heterochromatin formation and maintenance together with other proteins, including G9a [6, 7]. Some small-molecule inhibitors, including BIX-01294, UNC0638, and UNC0642, have been developed to specifically inhibit the catalytic activity of G9a/GLP, which provides a powerful tool to study this modification [8–10].

It has been well recognized that the heterochromatin is densely stained and distributed at the nuclear periphery based on the electron microscopy [11]. Through Lamin B1 (LMNB1) DamID technology, thousands of lamina-associated domains (LADs) have been identified in the mammalian genomes; these LADs dynamically approach nuclear lamina (NL) and contain fewer genes that are primarily repressed and late replicating [12, 13]. Interestingly, the genomic regions of large H3K9me2 domains (LOCKs) are highly overlapped with those of LADs [2]. Furthermore, via ^m6^A-tracer or fluorescent in situ hybridization (FISH)-based imaging, G9a-mediated H3K9me2 was proved to be required for chromatin-NL interactions [14, 15].

Besides large chromatin domains such as LADs, the genome is folded into multi-scale higher-order architecture in the eukaryotic nuclei. Each chromosome occupies its distinct territory in the nuclear space [16]. Based on chromatin conformation capture (3C)-based technologies, the 3D chromatin architecture has been revealed on different levels ranging about 100 kb to 10 Mb, including chromatin loops, topologically associating domains (TADs) and genomic compartments [17–19]. At the megabase scale, genomic compartments are segregated into two types, named A and B, that represent active and inactive chromatin domains, respectively [17]. Interestingly, B compartments are correlated with LADs [11, 20].

Based on the links among H3K9me2, LADs and B compartments, H3K9me2 could play roles in the 3D genome organization. In hematopoietic progenitors, it has been found that reduction of H3K9me2 by inhibiting G9a/GLP can change chromatin accessibility, as identified with the FAIRE technique [21]. However, details as to whether and how H3K9me2 may alter 3D chromatin architecture remain unknown. Hence, in the present study, we used Hi-C, DamID, ATAC-seq, and ChIP-seq technologies to study the functional roles of H3K9me2 in the 3D chromatin organization at the genome-wide scale.

## Results

### G9a/GLP inhibition removes H3K9me2 mainly at A compartments or inter-LADs in differentiated cells

To investigate the functional roles of H3K9me2 in the 3D genome organization, we used mouse AML12 hepatocytes as the model, as our previous studies showed that nuclear architecture were well preserved in AML12 cells [22, 23]. To remove H3K9me2, we treated AML12 cells with UNC0638, a selective inhibitor of G9a/GLP [9]. We observed a reduction of H3K9me2 levels by western blotting (WB) and immunofluorescence (IF) after UNC0638 treatment (**Figure S1A-B; Figure 1A**). It should be noted that the UNC0638 treatment cannot eradicate H3K9me2 completely, and some H3K9me2 signals remained at the nuclear periphery (**Figure 1A**). We also examined other modifications on the H3 lysine 9 residue and observed a slight decrease of H3K9me3 by WB (**Figure S1A-B**), which was similar to the result previously reported [24]. This observation may be on account of the “binary switch” on the same residue and non-processive H3K9 methylation mechanism by methyltransferases such as SUV39H1 [25, 26]. However, the nuclear distribution and genome-wide location of H3K9me3 and H3K9ac were hardly changed, as revealed by IF and ChIP-seq (**Figure S1C-D**). Additionally, we examined other covalent histone modifications (H3K27ac, H3K27me3, H3K4me1, and H3K4me3) and found no noticeable effect by WB and IF (**Figure S1E-F**).

**Figure 1.**
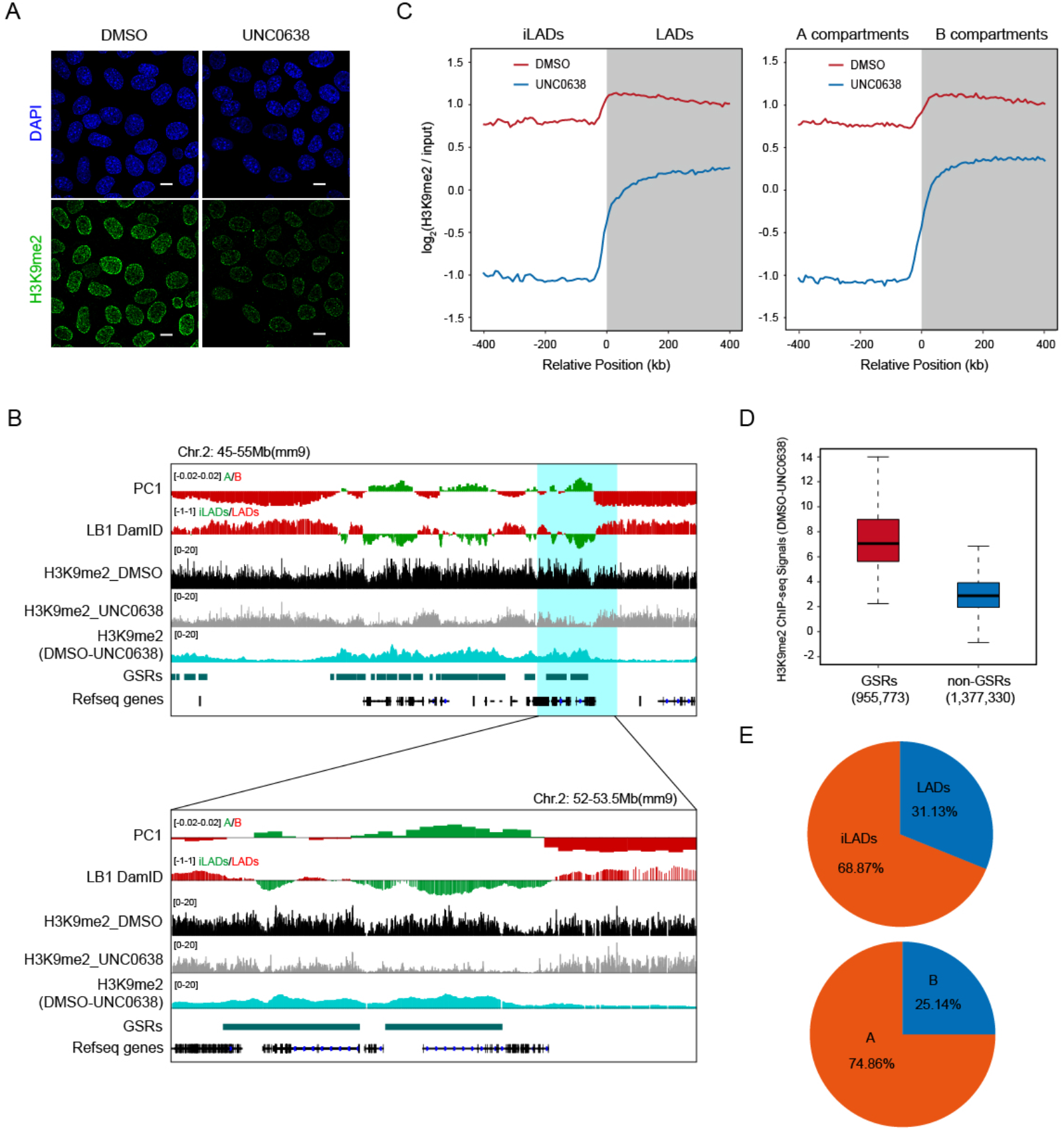
Region-specific removal of H3K9me2 modifications upon G9a/GLP inhibition. **A.** Representative IF images of H3K9me2 in DMSO and UNC0638 treated AML12 cells. Scale bar, 10 μm. **B.** Representative ChIP-seq tracks of H3K9me2 upon UNC0638 treatment in AML12 cells; PC1 values and smoothed LB1 DamID signals of AML12 cells [22]. Below: zoom-in view of the highlighted region. **C.** Aligned H3K9me2 profiles of mirrored border regions of A/B compartments (left) and LADs (right) in DMSO and UNC0638 treated AML12 cells. **D.** Box plots showing relative H3K9me2 levels (DMSO-UNC0638) at GSRs and non-GSRs in AML12 cells. The numbers at the bottom show the genomic bins of 1 kb. **E.** Percentages of GSRs in LADs/iLADs (upper) and A/B compartments (below) in AML12 cells.

Next, in order to investigate the profile of genome-wide H3K9me2 upon UNC0638 treatment in AML12 cells, we performed quantitative ChIP-seq experiments by spiking in human Hela cells. The degree of H3K9me2 reduction along the genome was not random as shown by ChIP-seq data, but it presented a region-specific pattern (**Figure 1B**). It should be noted that H3K9me2 ChIP-seq tracks showed a more or less equal distribution between LADs and inter-LADs (iLADs) when the data were not normalized. However, after normalizing the ChIP-seq data to input, the average H3K9me2 profile mainly decreased at iLADs or A compartments, but displayed a less degree of reduction at LADs or B compartments (**Figure 1C**). This data was in agreement with the observation by IF imaging.

Next, we called subcompartments in AML12 cells based on a reported method [19], by integrating Hi-C contact matrices, histone modifications and lamina-association maps (**Figure S1G-H**). We then examined H3K9me2 reduction in these subcompartments after UNC0638 treatment. The degree of H3K9me2 reduction in A1 and A2 subcompartments were higher than those in B subcompartments. Moreover, the B2 subcompartments, which are with the higher H3K9me3 level, are more resistant to UNC0638 treatment than B1 and B3 compartments (**Figure S1I)**.

Consistently, a reanalysis of published ChIP-seq data of G9a knockout and wild-type MEF cells [27] also showed a specific reduction of H3K9me2 at iLADs (**Figure S2A**). Furthermore, G9a depletion in mouse cardiomyocytes *in vivo* resulted in similar changes in H3K9me2 patterns [28] (**Figure S2B**). It should be noted that H3K9me2 were “increase” at LADs in G9a/GLP knockout samples, because these datasets were not generated by using spike-in method and thus could be not quantitative. These results indicate that neither chemical nor genetic inhibition of G9a/GLP completely removes H3K9me2 modification at LADs/B compartments.

Considering the different removal pattern of H3K9me2 after G9a/GLP inhibition, we then referred these dramatically H3K9me2-decreased chromatin regions as G9a/GLP-sensitive regions (GSRs) (**Figure 1D**), which occupied 37.5% of the genomic regions in AML12 cells (**Table S1**). There are 68.87% and 74.86% of GSRs locating within iLADs and A compartments in AML12 cells, respectively (**Figure 1E**). These results suggest that the sensitivity of H3K9me2-modified regions upon G9a/GLP inhibition demarcates two different types of genomic compartments, which are analogous to A/B compartments or iLADs/LADs.

### G9a/GLP inhibition removes H3K9me2 globally in mESCs

As H3K9me2 modifications are highly dynamic during stem cell differentiation [2, 10], we wondered whether the sensitivity of G9a/GLP inhibition on H3K9me2 was cell-type specific. Treatment of mouse embryonic stem cells (mESCs) with UNC0638 induced a decrease of H3K9me2 (**Figure S3A**). Unlike the results in AML12 cells, H3K9me2 at both the nuclear interior and periphery were almost removed completely, and there was a foci-like staining of H3K9me2 in mESCs treated with UNC0638 as shown by IF images, which was similar to H3K9me3 staining (**Figure S3B**). A similar result was also found in G9a^−/−^ mESCs [29]. Consistently, ChIP-seq data also showed that most of H3K9me2 modification, including those in LADs/B compartments, were removed entirely, but some “sharp peaks” remained (**Figure S3C-D**). These retained H3K9me2 peaks were mainly (70.3%) overlapped with H3K9me3 peaks (**Figure S3E**). We also analyzed the features of the remaining non-overlapping peaks, and found that they were mostly intergenic and enriched for repeat sequences (**Figure S3F-G**). In addition, the ChIP-seq data of GLP-knockout mESCs [27] yielded similar results (**Figure S3C**). Therefore, unlike differentiated cells such as AML12 and MEFs, H3K9me2 modification in mESCs are mainly G9a/GLP-sensitive. Although H3K9me2 were globally removed after G9a/GLP inhibition in mESCs, the A/B compartments were largely maintained (**Figure S3H**).

### H3K9me2 insensitive to G9a/GLP inhibition is associated with genomic compartments

As shown by the representative region, the tracks of LB1 DamID and PC1 displayed similar wavy patterns in AML12 cells; after UNC0638 treatment, H3K9me2 was more reduced at regions with weaker LB1 DamID signals and stronger A-tendency PC1 values, in both iLADs/A and LADs/B, and the wavy patterns of remaining H3K9me2 were more similar to those of LB1 DamID and PC1 tracks (**Figure 2A**). Quantitative analysis showed that, the H3K9me2 levels after G9a/GLP inhibition exhibited much stronger correlation with PC1 values (r = −0.661) or LB1 DamID signals (r = 0.701), comparing with the DMSO-treated samples (r = −0.198 for PC1, r = 0.205 for LB1 DamID; **Figure 2B-C**). However, other inactive histone modifications such as H3K9me3 and H3K27me3, displayed little correlation with PC1 or LB1 DamID (**Figure 2D-E**).

**Figure 2.**
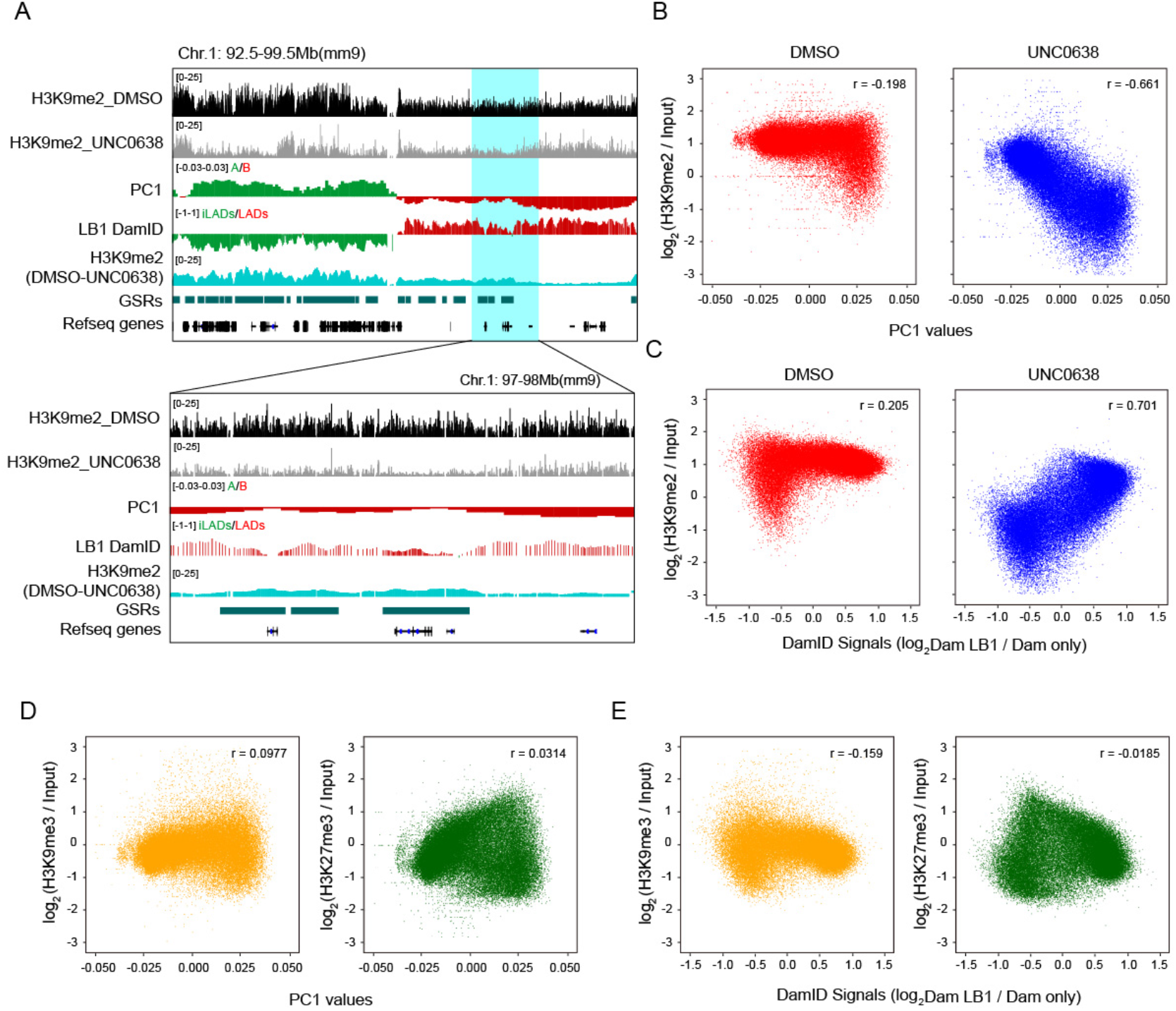
G9a/GLP-insensitive H3K9me2 is associated with compartmental status. **A.** Representative tracks of H3K9me2 ChIP-seq and difference (DMSO-UNC0638) in DMSO and UNC0638 treated AML12 cells; PC1 values and smoothed LB1 DamID signals of AML12 cells [22]. Below: zoom-in view of the highlighted region. **B.** Correlation between H3K9me2 levels and PC1 values in DMSO (left) and UNC0638 (right) treated AML12 cells. **C.** Correlation between H3K9me2 levels and LB1 DamID signals in DMSO (left) and UNC0638 (right) treated AML12 cells. **D.** Correlation between PC1 values and H3K9me3 (left) / H3K27me3 (right) levels in AML12 cells. **E.** Correlation between LB1 DamID signals and H3K9me3 (left) / H3K27me3 (right) levels in AML12 cells. The 40 kb bins were used for the correlation analyses shown from C to F.

Histone modifications can reflect different chromatin state segments, which are associated with fine-scale genomic compartmentalization [30]. Our data further suggest that the intrinsic H3K9me2 modification at non-GSRs could better reflect the inactive environment of chromatin and are highly correlated with the genomic compartments.

### G9a/GLP inhibition decreases chromatin-NL interactions primarily at GSRs

Our electron microscopy data showed that less nuclear periphery heterochromatin was seen in UNC0638 treated AML12 cells (**Figure 3A**). To investigate the effects of G9a/GLP inhibition on chromatin-NL interactions at a genome-wide scale, we conducted LB1 DamID assay in AML12 cells. By examining the DamID signals, the loss or gain of LADs occurred in some genomic regions (**Figure 3B**), and the coverage of LADs decreased from 54.75% to 47.43% (**Figure 3C; Table S2**). These switched LADs or iLADs are associated with the significant changes of gene expression levels (**Figure 3D**).

**Figure 3.**
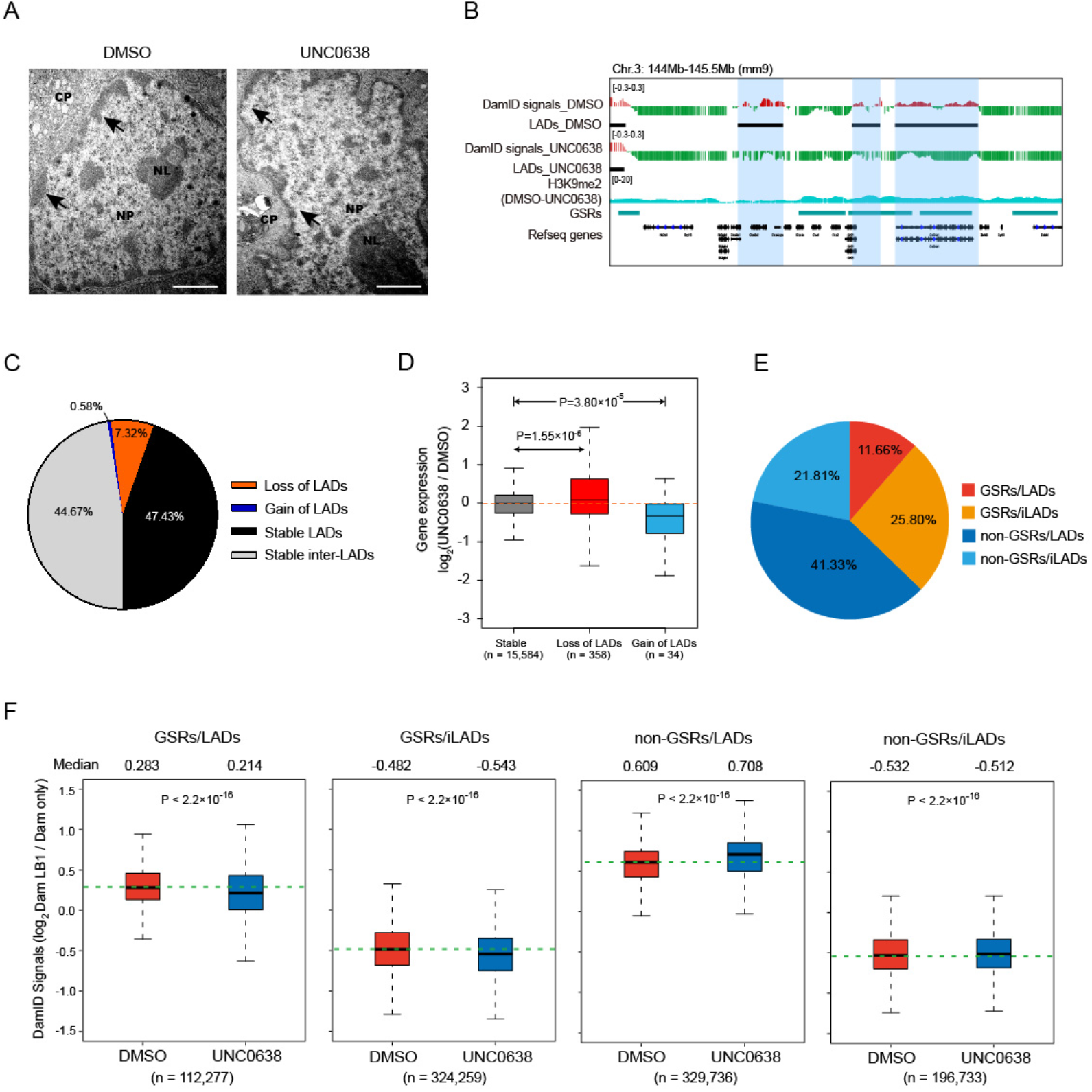
Inhibition of G9a/GLP by UNC0638 treatment decreases chromatin-NL interactions mainly at GSRs. **A.** Representative electron micrographs of DMSO and UNC0638 treated AML12 cells. CP, cytoplasm; NP, nucleoplasm; NL, nucleolus. Black arrowheads indicate the nuclear periphery chromatin. Scale bar, 2 μm. **B.** Representative tracks of smoothed LB1 DamID signals and H3K9me2 ChIP-seq data of DMSO and UNC0638 treated AML12 cells. Black bars represent locations of LADs, and colored shadows indicate switched regions. **C.** Proportions of switched and stable LADs in AML12 cells upon UNC0638 treatment. **D.** Box plots showing relative expression changes of genes associated with loss or gain of LADs in AML12 cells upon UNC0638 treatment. P-values, Wilcoxon rank-sum test. **E.** Proportions of the four sections (GSRs/LADs, GSRs/iLADs, non-GSRs/LADs and non-GSRs/iLADs) in the genome of AML12 cells. **F.** Box pots of LB1 DamID signals of the four sections in DMSO and UNC0638 treated AML12 cells. The numbers at the bottom show the probes counts of LB1 DamID. P-values, *t*-test.

In order to further study the connection between H3K9me2 and chromatin-NL interactions, we classified the genome into four sections: GSRs/LADs (11.66%), GSRs/iLADs (25.8%), non-GSRs/LADs (41.33%) and non-GSRs/iLADs (21.81%) (**Figure 3E**). Because of the large sample size in the statistical tests, chromatin-NL interactions are significantly different in all these sections, but the degree and direction of changes are variable. As shown in **Figure 3F**, the chromatin-NL interactions in GSRs/iLADs and GSRs/LADs decreased after UNC0638 treatment, consistent with the obvious decrease of H3K9me2 in these regions. However, in non-GSRs, where H3K9me2 were less reduced, the chromatin-NL interactions increased (non-GSRs/LADs) or were hardly changed (non-GSRs/iLADs). Of note, the detected increase of DamID signal in non-GSRs/LADs could be not biologically relevant, but instead resulted from the normalization of DamID data of UNC0638-treated samples, in which chromatin-NL interactions were reduced in GSRs. Therefore, G9a/GLP inhibition decreases chromatin-NL interactions mainly at GSRs.

### G9a/GLP inhibition increases genomic compartmentalization

To further investigate the impacts of H3K9me2 on the 3D genome organization, we generated high-resolution chromatin interaction maps by *in situ* Hi-C assay in DMSO and UNC0638-treated AML12 cells. We performed two biological replicates for the Hi-C experiments and obtained highly consistent results (**Figure S4A-C**). Then, by overviewing the long-range Hi-C contact maps, we can see interaction changes among compartments (**Figure 4A**). Further analysis of genomic compartmentalization by pairwise comparison of chromatin interactions between ranked PC1 intervals revealed that the interactions between A and B compartments decreased, while the interactions among the same type of compartments (A vs. A; B vs. B) increased (**Figure 4B; Figure S4D**). Through quantification with the duplicates of Hi-C data, we showed that the compartmentalization strength [31], increased by 23%, from 3.5 to 4.3, upon G9a/GLP inhibition (**Figure 4C**). Similarly, the strength of TAD boundaries between A and B compartments significantly increased, but those within A or B compartments were not obviously changed, which indicated the increased insulation between A/B compartments (**Figure 4D; Table S3**). These results suggest the increased degree of genomic compartmentalization upon G9a/GLP inhibition, coincident with the enlarged difference of H3K9me2 levels between A and B compartments.

**Figure 4.**
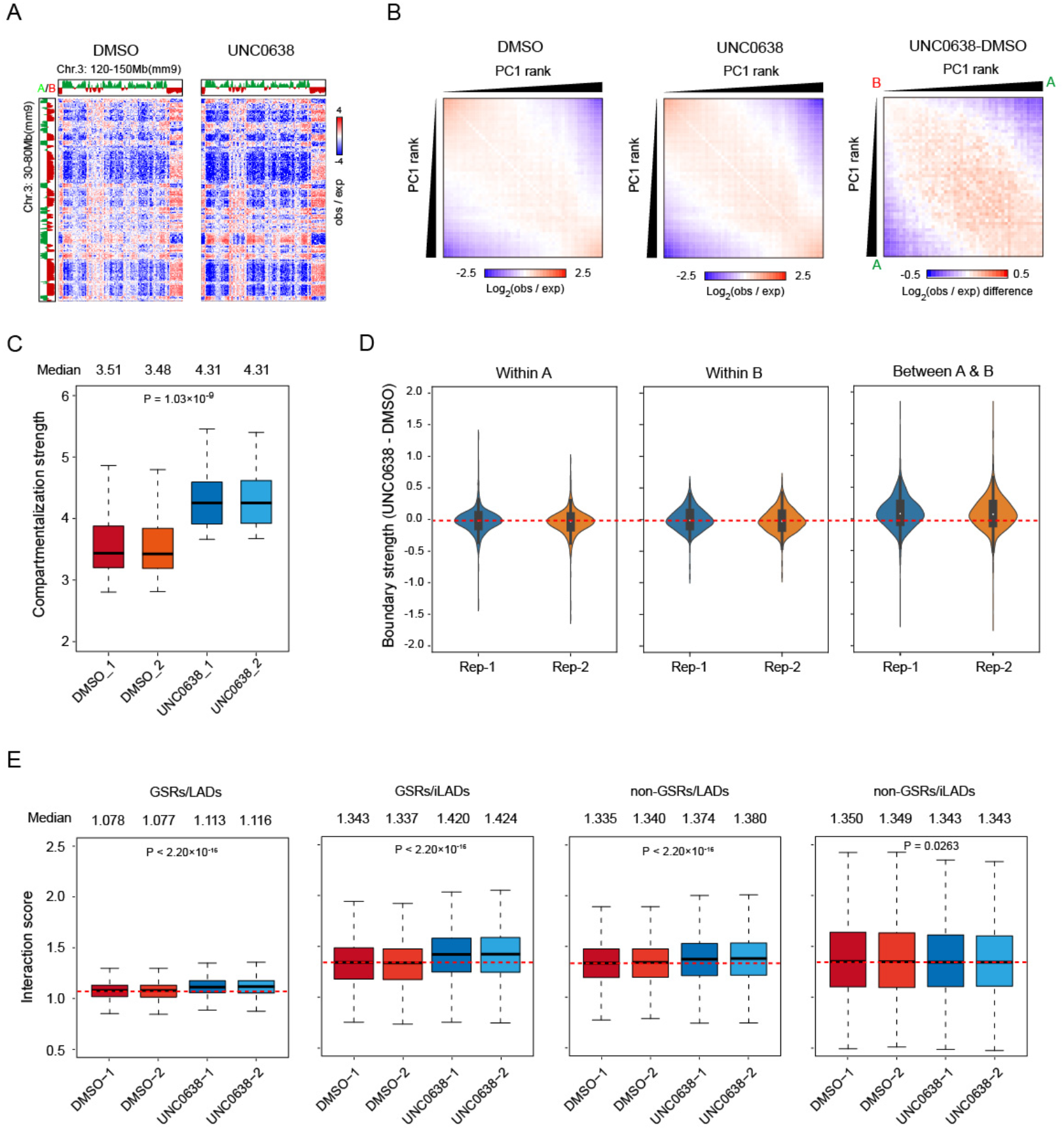
Removal of H3K9me2 at GSRs increases genomic compartmentalization. **A.** Representative Hi-C contact matrices (obs/exp) in DMSO (left) and UNC0638 (right) treated AML12 cells. **B.** Average contact enrichment between pairs of 250 kb bins ranked by PC1 values in DMSO (left) and UNC0638 (middle) treated AML12 cells, and the difference between them (right). **C.** Box plots showing compartmentalization strength across chromosomes in DMSO and UNC0638 treated AML12 cells. P-value, two-factor ANOVA. **D.** Violin plots showing TAD boundary strength changes within A (left), B (middle) and between A/B (right) in AML12 cells upon UNC0638 treatment. **E.** Box plots showing interaction scores of the four fractions (GSRs/LADs, GSRs/iLADs, non-GSRs/LADs and non-GSRs/iLADs) in DMSO and UNC0638 treated AML12 cells. P-values, two-factor ANOVA.

Furthermore, via the stratification of the genome by considering GSRs and LADs, we found that interaction scores, which measure the chromatin interactions at given regions [23], increased significantly in all sections except for non-GSRs/iLADs after UNC0638 treatment, consistent with the results of chromatin-NL interactions (**Figure 4E**). These results are in agreement with the previous report that chromatin-NL are coupled with chromatin-chromatin interactions in the genome organization [32]. In non-GSRs/LADs where H3K9me2 is less removed, the increased chromatin-chromatin interactions could be the secondary effect of chromatin changes at GSRs after H3K9me2 reduction. Similarly, it has been reported that active chromatin marks on euchromatin can drive spatial sequestration of heterochromatin indirectly in differentiated cells [33].

### H3K9me2 reduction by G9a/GLP inhibition alters the expression of hundreds of genes

To further investigate the biological functions of G9a/GLP-sensitive H3K9me2 in AML12 cells, we treated the cells with UNC0638 and examined gene expression by RNA-seq. There were 709 genes with significant expression changes upon UNC0638 treatment (**Table S4**). Among them, 483 genes (68%) were up-regulated, and 226 genes (32%) were down-regulated (**Figure 5A**). Gene Ontology (GO) analysis showed that up-regulated genes were mainly enriched in liver biological processes, including steroid biosynthesis and lipid metabolism (**Figure 5B**), yet down-regulated genes had no significant enrichment.

**Figure 5.**
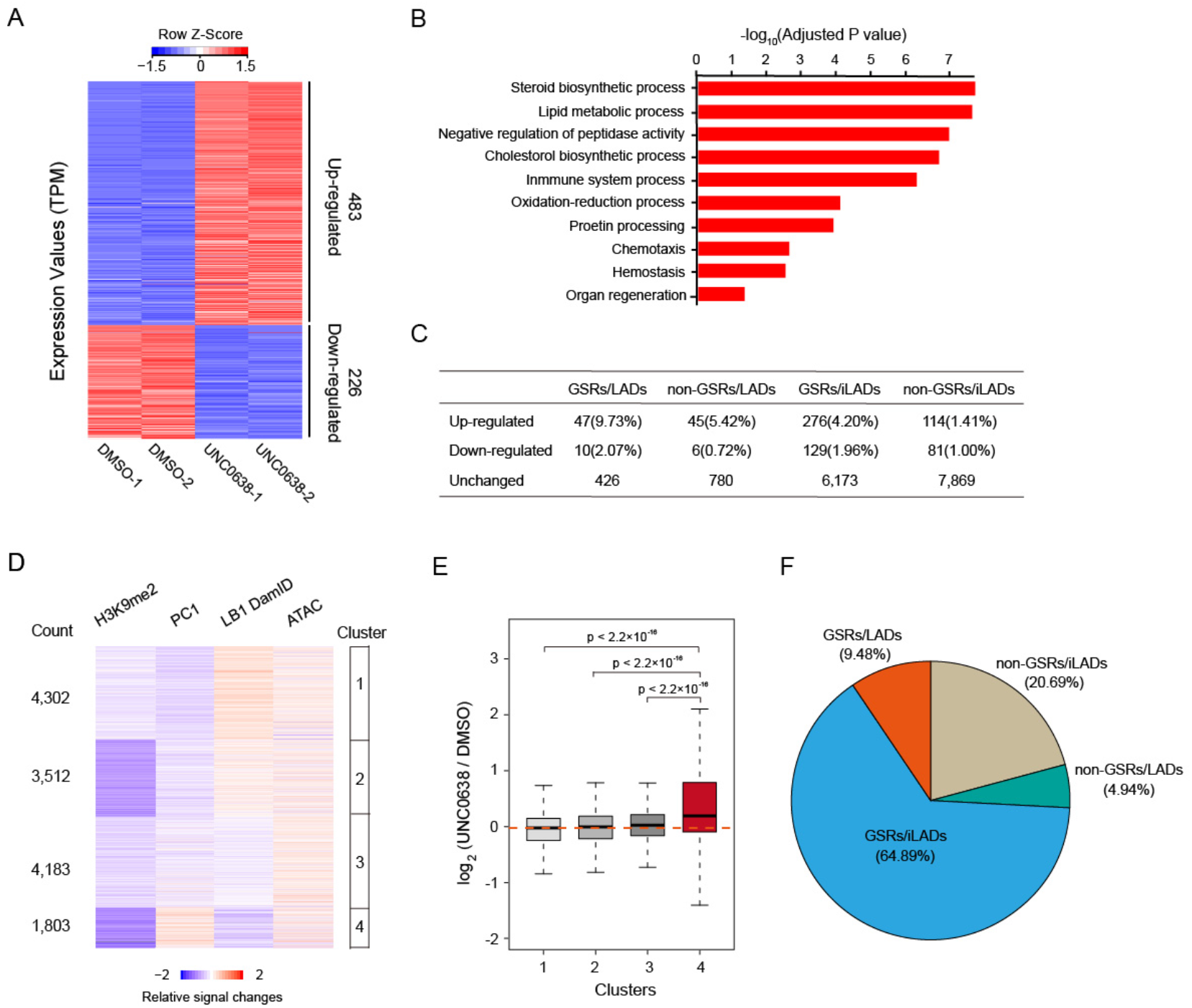
Gene expression changes upon UNC0638 treatment in AML12 cells. **A.** Heatmap of differentially expressed genes between DMSO and UNC0638 treated AML12 cells, with two biological replicates (independent DMSO/UNC0638 treatment, RNA libraries preparation and sequencing) for each sample. **B.** Top ten Gene Ontology (GO) terms of up-regulated genes upon UNC0638 treatment in AML12 cells. **C.** Quantification of expression changed/unchanged genes in four fractions (GSRs/LADs, non-GSRs/LADs, GSRs/iLADs and non-GSRs/iLADs) in AML12 cells. The percentages of up-regulated genes in each fractions were shown. **D.** Clusters of gene-centric signal changes of H3K9me2 ChIP-seq signals, PC1 values, DamID signals and ATAC-seq signals upon UNC0638 treatment in AML12 cells. Relative signal changes were calculated at promoters with bin size of 5 kb. **E.** Box plots showing gene expression changes in four clusters upon UNC0638 treatment in AML12 cells. **F.** Proportions of the cluster 4 genes in four sections (GSRs/LADs, GSRs/iLADs, non-GSRs/LADs and non-GSRs/iLADs) in AML12 cells.

To explore the relationship between gene expression and chromatin organization, we examined genes in the context of GSRs and LADs. In all the four sections (GSRs/LADs, GSRs/iLADs, non-GSRs/LADs, non-GSRs/iLADs), the numbers of up-regulated genes were more than those of down-regulated genes (**Figure 5C**). Usually, the gene up-regulation is associated with the increased chromatin accessibility. Thus, we performed ATAC-seq to examine the chromatin accessibility changes upon UNC0638 treatment and combined with the high-throughput data, including H3K9me2 ChIP-seq, RNA-seq, DamID and Hi-C, for an integrated analysis. After excluding the genes that were not expressed in both DMSO and UNC0638-treated samples, we used a gene-centric approach to aggregate relative signal changes of these chromatin features and identified 4 clusters showing different chromatin alterations (**Figure 5D; Table S5**). In all the 4 clusters, H3K9me2 more or less decreased and the ATAC-seq signals increased averagely, indicating the increased chromatin accessibility at promoters of these genes. Among them, cluster 4 shows decreased H3K9me2, increased Hi-C PC1 (compartment A tendency) and decreased chromatin-NL interactions (DamID) (**Figure 5D**). Genes in cluster 4 tended to be up-regulated after UNC0638 treatment; however, the gene expression levels of other three clusters were not obviously affected (**Figure 5E**). Moreover, genes in cluster 4 were mainly enriched in GSRs (64.89% in GSRs/iLADs, 9.48% in GSRs/LADs; **Figure 5F**). These results indicate that the removal of G9a/GLP-sensitive H3K9me2 up-regulates the expression of hundreds of genes, which are associated with the more open chromatin environment.

## Discussion

In this study, we show that G9a/GLP inhibition mainly removes H3K9me2 at A compartments, and G9a/GLP-insensitive H3K9me2 is highly correlative with genomic compartments globally, suggesting that the level of G9a/GLP-sensitivity of H3K9me2 could reflect the intrinsic tendency of chromosomes to segregate into diverse compartments. Chromatin compartmentalization can be regulated by histone modifications through phase separation [34]. In this study, we show that the removal of G9a/GLP-sensitive H3K9me2 in AML12 cells increases the genomic compartmentalization and boundary strength of TADs between A and B compartments, and decreases chromatin-NL interactions heterogeneously that are associated with H3K9me2 reduction. Additionally, H3K9me2 reduction by G9a/GLP inhibition up-regulates hundreds of genes associated with alterations of chromatin organization. Our results thus revealed the functional roles of H3K9me2 in the 3D chromatin organization. It has been shown that, based on immunostaining data, G9a is mainly responsible for H3K9 modifications at euchromatic loci (i.e., nuclear areas other than the DAPI-dense pericentromeric heterochromatin) in mESCs [29]. In this article, we further demonstrated that regardless of chemical inhibition or genetic depletion of G9a/GLP, H3K9me2 was not removed completely in B compartments or LADs in differentiated cells. However, H3K9me2 can be more effectively removed in mESCs by G9a/GLP inhibition. Moreover, the induction of mESCs to epiblast-like cells (EpiLCs) and primordial germ cell-like cells (PGCLCs) is accompanied by a large-scale reorganization of chromatin signatures, including H3K9me2 patterns [35]. During hematopoietic stem cell lineage commitment, G9a/GLP-dependent H3K9me2 marks are established gradually [10]. These data imply a complexity of mechanisms underlying H3K9me2 modification in differentiated cells. The incomplete removal of H3K9me2 by G9a/GLP inhibition in differentiated cells could imply an inefficient UNC0638 treatment at B compartments or a G9a/GLP-independent H3K9me2 modifying system existing in differentiated cells. A recent study reported that the histone turnover of heterochromatin at nuclear periphery are repressed, which may also explain the insensitivity of H3K9me2 to G9a/GLP [36].

However, it remains unclear what chromatin modifiers could be responsible for the remaining H3K9me2 modifications at inactive compartments of differentiated cells after G9a/GLP inhibition. The Suv39 family members, including SUV39H1/2 and SETDB1/2, are also known to be involved in the H3K9me2 modification [1]. It has been reported that during somatic cell reprogramming into iPSCs, H3K9 methylation is a crucial barrier and knockdown of *Suv39h1/2* and *Setdb1* can promote the reprogramming, but the H3K9me2 level was not significantly decreased at some gene loci by ChIP-qPCR examination [37]. A recent report also showed that even triple-knockout of *Suv39h1/2* and *Setdb1* did not cause a marked decrease of H3K9me2 in mouse liver [38]. It would be fascinating to further investigate the modifiers of these G9a/GLP-insensitive H3K9me2 marks and their functions in chromatin organization and other biological processes in the future.

The eukaryotic nuclear lamina is mainly composed of Lamin A/C and Lamin B, as well as other interacting components inside or nearby the networks, which provides a depressing environment for LADs. G9a was found to interact with the NL-interacting protein BAF [39]. Also, G9a and GLP play novel roles in regulating heterochromatin anchorage to the nuclear periphery via the methylation and stabilization of Lamin B1, which associates with H3K9me2-marked peripheral heterochromatin [40]. H3K9me3 in LADs was reported to be combined with lamin B receptor (LBR) via mediation by CBX5 [41]. In mammalian cells, the histone deacetylase, HDAC3, was reported to interact with the Lamin-associated protein LAP2β to maintain the repressed state of peripheral chromatin [42]. In the current study, we provided genomic data showing that G9a/GLP inhibition mainly decreases chromatin-NL interactions of LADs at GSRs, whereas LADs at non-GSRs remain intact. Therefore, for a better understanding of the organization and functions of LADs not affected by G9a/GLP inhibition, further efforts in the field would be needed to decipher factors associated with G9a/GLP-insensitive H3K9me2 at the nuclear periphery.

## Materials and methods

### Cell culture and UNC0638 treatment

The mouse hepatocyte cell line alpha mouse liver 12 (AML12, ATCC^®^ CRL-2254™) was cultured in 37 °C and 5% CO_2_ incubator in medium DMEM/F12 (ThermoFisher, 11320033, Waltham, MA, USA) supplemented with 10% fetal bovine serum (FBS, Gibco, 16140071), 1× ITS Liquid Media Supplement (100×, Gibco, 41400045) and 40ng/ml dexamethasone (Sigma, D4902, Darmstadt, Germany). AML12 cells were treated with 8 μM of UNC0638 (Selleck, S8071, Houston, TX, USA) for five days, and the same amount of DMSO (Sigma, D2650) was used as the control.

Mouse embryonic stem cell line E14 (E14TG2a) was cultured in 2i/LIF conditions as described [12]. The G9a/GLP inhibitor UNC0638 was added into the medium to a final concentration of 0.5 μM for five days of treatment, and the same amount DMSO as control.

### Western blotting

Cells lysis was incubated at 95℃ for 15min with 1×SDS-PAGE Loading Buffer. The primary antibodies are as follows: anti-H3 (Abcam, ab1791), anti-H3K9me2 (Abclonal, A2359), anti-H3K9me3 (Abclonal, A2360), anti-H3K9ac (Active Motif, 39137), anti-H3K27ac (Active Motif, 39133), anti-H3K27me3 (Cell Signaling Technology, 9733), anti-H3K4me1 (Active Motif, 39297) and anti-H3K4me3 (Abclonal, A2357). HRP-conjunction secondary antibodies (Jackson ImmunoResearch Laboratories, PA, USA) and Chemiluminescent HRP Substrate (Millipore, WBKLS0500) were used for the detection. Software AlphaView was used for the relative grayscale statistics.

### Immunofluorescence (IF) Assay

Cells were fixed in 4% formaldehyde followed by the treatment with 0.5% Triton X-100 for 10min at room temperature. The cells were blocked with phosphate buffered saline (PBS) containing 4% bovine serum albumin (BSA) for 30min at room temperature and then processed for immunostaining. The primary antibodies are same as that we used for western blotting and the secondary antibodies are as follows: Alexa Fluor 488-AffiniPure Donkey Anti-Rabbit IgG (Jackson ImmunoResearch Laboratories), Alexa Fluor 594-AffiniPure Donkey Anti-Rabbit IgG (Jackson ImmunoResearch Laboratories). The slides were counterstained with DAPI (Beyotime, C1002). Fluorescence images were taken with Leica confocal microscope (TCS SP5, Leica, Germany) and we controlled the confocal parameters unchanged in each set of experiments. The images were analyzed by the LAS AF Lite software.

### Transmission electron microscopy

Cells were scratched and collected followed by fixed in 2.5% glutaraldehyde (Sigma) diluted in 0.1M Phosphate Buffer (PB, pH7.4). The samples were further processed with 1% Osmium tetroxide, dehydrated with acetone and embedded with resin, at the EM facility in the School of Basic Medical Sciences, Fudan University. Target cells were randomly selected and captured with a transmission electron microscope (JEOL-1230, Nippon Tekno, Japan).

### ChIP-seq

The ChIP experiments were performed as described [12] with the antibodies anti-H3K9me2 (Abcam, ab1220), anti-H3K9me3 (Abcam, ab8898), H3K9ac (Active motif, 39137) and anti-CTCF (CST, 3418). To calibrate H3K9me2 ChIP-seq, Hela cells were mixed with DMSO or UNC0638 treated AML12 cells in 1:4 proportion initially. The libraries were prepared using the NEBNext Ultra II DNA Library Prep Kit (NEB) followed by NG sequencing using the Illumina HiSeq X Ten system. Two biological replicates (independent DMSO/UNC0638 treatment, ChIP assay and sequencing) were performed. Raw sequencing reads were mapped to the mouse mm9 genome reference using Bowtie2 [43]. Duplicated read pairs were discarded using Samtools [44]. log_2_ ratio of IP and input signals were calculated using deepTools [45]. For the normalization of spike-in H3K9me2 ChIP-seq data, the NGS data were mapped to refseq genome of mm9 (AML12) and hg19 (Hela), respectively. Then, the Input, DMSO-treated and UNC0638-treated AML12 H3K9me2 ChIP-seq signals were divided by the ratio of unique mapping reads [human / (human + mouse)].

To define G9a/GLP-sensitive regions (GSRs), we firstly subtracted DMSO and UNC0638 treated ChIP signals with bins of 1kb. Then, the data was normalized and smoothed using a moving average approach with 40kb window size. At each bin, we converted the smoothed signals into the t-statistics and then determined a threshold −8 for calling domains based on probe level FDR 0.01 using the left part of the distribution as null. GSRs were defined as regions with consecutive bins with t-statistics greater than the threshold, and the length of domains was limited to no less than 50 kb. Meanwhile, domains with distances less than 10 kb were merged.

### RNA-seq

RNA was extracted with TRIzol Reagent (Life technology, 15596018). The RNA libraries were prepared using the Ribo-Zero Gold rRNA Removal Reagent (Illumina) and NEBNext Ultra II RNA Directional Library Prep Kit (NEB) followed by NG sequencing using the Illumina HiSeq X Ten system. Two biological replicates (independent DMSO/UNC0638 treatment and sequencing) were performed. Paired-end sequencing reads were mapped to the mouse mm10 genome reference by HISAT2 [46]. To identify differentially expressed genes, reads mapped to annotated genes were counted with the HTSeq package [47]. Fold-change and p values were calculated by DESeq2 [48]. FDR < 0.05 and |log_2_(fold change)| > 1 were used as thresholds. GO terms enrichment analysis was performed using the DAVID tool [49].

To define gene-centric changes of chromatin features, we removed genes that were not expressed in both DMSO and UNC0638 treated samples (gene baseMean less than 10.30 in DESeq2) and calculated relative signal changes at promoters with bin size of 5 kb. We used difference (UNC0638 - DMSO) for PC1 values and DamID signals, and ratio (log_2_(UNC0638 / DMSO)) for signal changes of ATAC-seq and ChIP-seq. Then we used K-means clustering to divide genes into 4 categories with different chromatin feature changes.

### DamID

Lamin B1 (LB1) DamID experiments were performed as described [12]. Microarray hybridization assay and data analysis were as described [50]. Two biological replicates (independent DMSO/UNC0638 treatment, DamID and microarray hybridization assay) were performed.

### ATAC-seq

ATAC experiments and data analysis were conducted as described [51] with minor modifications: harvesting 30,000 AML12 cells followed by transposition reaction at 37℃ for 45 min. Libraries were sequenced via the Illumina HiSeq X Ten system. Two biological replicates (independent DMSO/UNC0638 treatment, ATAC assay and sequencing) were performed. Paired-end reads were mapped to the mouse genome (mm9) using Bowtie2 with parameters “-X 2000”. Mitochondrial and PCR duplicate reads were discarded after alignment. Peaks calling was used MACS2 [52] with a q value threshold of 0.01.

### *In situ* Hi-C

The *in situ* Hi-C libraries were prepared as previously described [19]. The libraries were then sequenced via the Illumina HiSeq X Ten system. Two biological replicates were performed (independent DMSO/UNC0638 treatment, Hi-C assay and sequencing). Paired-end reads were aligned to mm9 reference genome using HiC-Pro [53].

To call compartments, eigenvector function from juicer [54] was used to calculate PC1 values at 40 kb resolution. The sign of PC1 values was adjusted based on gene density to assign A and B compartments.

To call TADs, HiCtool [55] based on the directionality index and hidden Markov model was used at 40kb resolution. To compare the structure of TADs between two conditions, “the distance between the centers of two boundaries less than or equal to 40kb” was used as criteria. To compare TAD boundary strength, the script matrix2insulation.pl [56] was used from cworld::dekker.

To compare interactions between two conditions, the ratio of observed and expected normalized counts was calculated by dump function from juicer.

To calculate compartmentalization strength, we used median (AA, BB) / median (AB) as a measure, where AA is obs/exp between pairs of loci with a strong A compartment (top 20% based on PC1 values) signal, BB is obs/exp between pairs of loci with a strong B compartment (bottom 20% based on PC1 values) signal, and AB is obs/exp between pairs of loci with these strong A and B compartment signal.

To calculate interaction scores, we analyzed them at the bin level of 40kb resolution according to its PC1 values. For each bin, we calculated its average interaction frequencies with bins of the same section (C_x_, the same section means GSRs/LADs, GSRs/iLADs, non-GSRs/LADs or non-GSRs/iLADs) and its average interaction with any other bins (C_total_) using the obs/exp. We calculated the interaction score as CS_x_ = C_x_ / C_total_.

## Data access

All the data of DamID, ATAC-seq, RNA-seq, ChIP-seq, and Hi-C from this study are submitting to GSA at the National Genomics Data Center (https://bigd.big.ac.cn/) under accession number CRA002762. A link to a UCSC genome browser session displaying the uploaded sequence tracks: https://genome.ucsc.edu/s/Wenlab/H3K9me2_updated_tracks

## Authors’ contributions

ZY, LJ and BW conceived and designed this study. ZY and XH performed most of experiments; QW prepared Hi-C libraries; YZ performed certain western blot assays. LJ performed the bioinformatics analyses. ZY, XH, LJ and BW prepared the manuscript.

## Acknowledgements

We thank members of the Wen lab for their suggestions and supports, J. Li and Y. Huang for assistance with confocal microscope operation, S. Dong from F. Zhang Lab for assistance with microarray assay of DamID. This work was supported by the National Key Research and Development Program of China (2018YFC1003500 to B.W.), and the National Natural Science Foundation of China (31771435 to B.W.).

## Supplementary material

**Figure S1.**
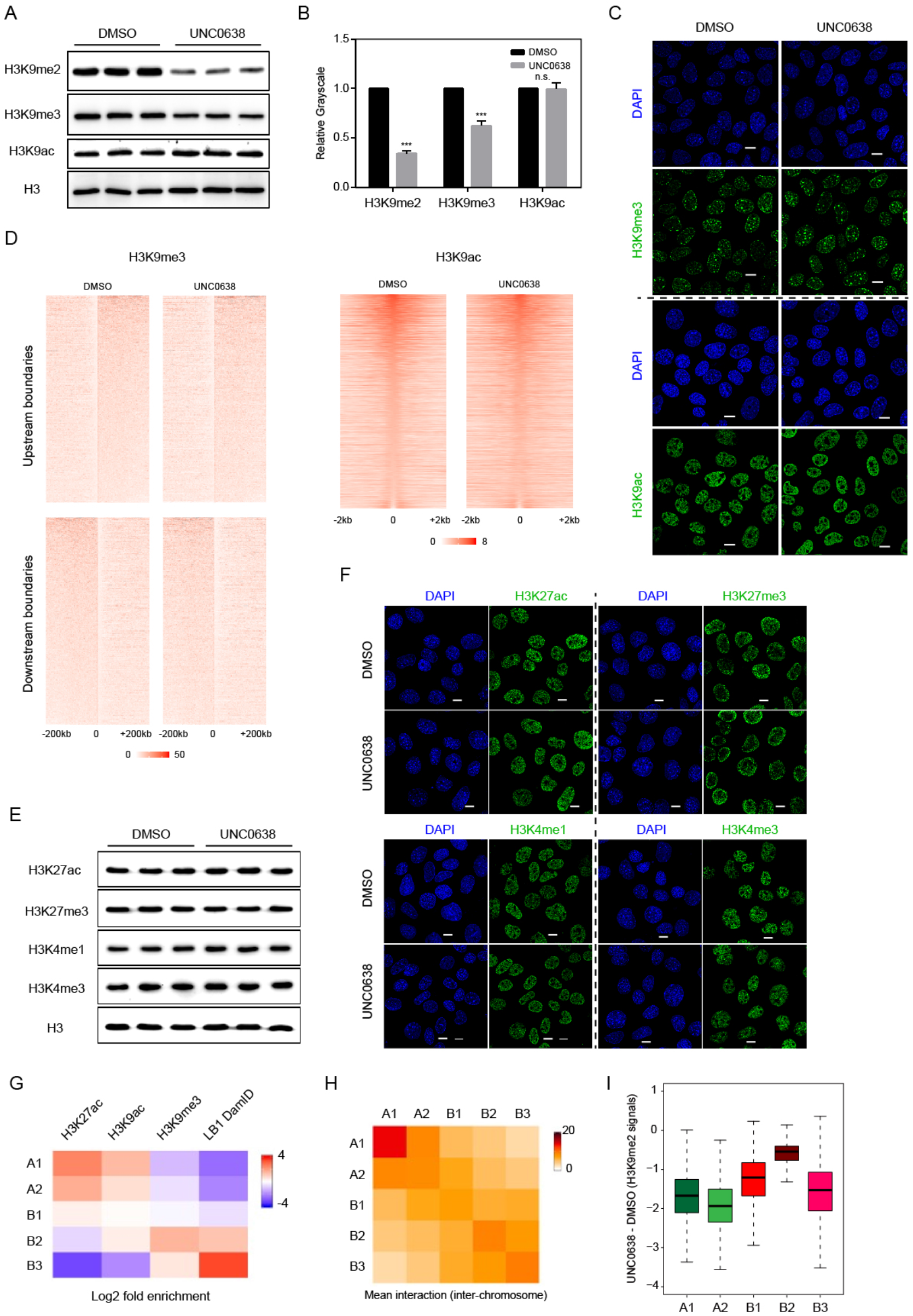
Global levels of histone modifications upon UNC0638 treatment in AML12 cells. **A.** Western blotting with antibodies against H3K9me2, H3K9me3 and H3K9ac in DMSO and UNC0638 treated AML12 cells, with three biological repeats respectively (independent DMSO/UNC0638 treatment). H3 serves as the loading control. **B.** Relative grayscale of western blotting from (A) calculated by AlphaView software. P-values, *t*-test. ***, P < 0.001. **C.** Representative IF images of H3K9me3 and H3K9ac in DMSO and UNC0638 treated AML12 cells. Scale bar, 10 μm. **D.** Genome-wide alignments of H3K9me3 (left) and H3K9ac (right) ChIP-seq in DMSO and UNC0638 treated AML12 cells. The center of H3K9me3 heatmap is the boundary of its domain, and the center of H3K9ac heatmap is the called peak center. **E.** Western blotting with antibodies against H3K27ac, H3K27me3, H3K4me1 and H3K4me3 in DMSO and UNC0638 treated AML12 cells, with three biological repeats respectively (independent DMSO/UNC0638 treatment). H3 serves as the loading control. **F.** Representative IF images of H3K27ac, H3K27me3, H3K4me1 and H3K4me3 in DMSO and UNC0638 treated AML12 cells. Scale bar, 10 μm. **G.** Heatmap showing the log_2_ fold enrichment of H3K27ac、H3K9ac、H3K9me3 and LB1 among five different subcompartments. **H.** Contact enrichment among the five subcompartments. Mean Interaction between five different compartments. **I.** Box plots showing the relative H3K9me2 levels (DMSO-UNC0638) at subcompartments after UNC0638 treatment in AML12 cells.

**Figure S2.**
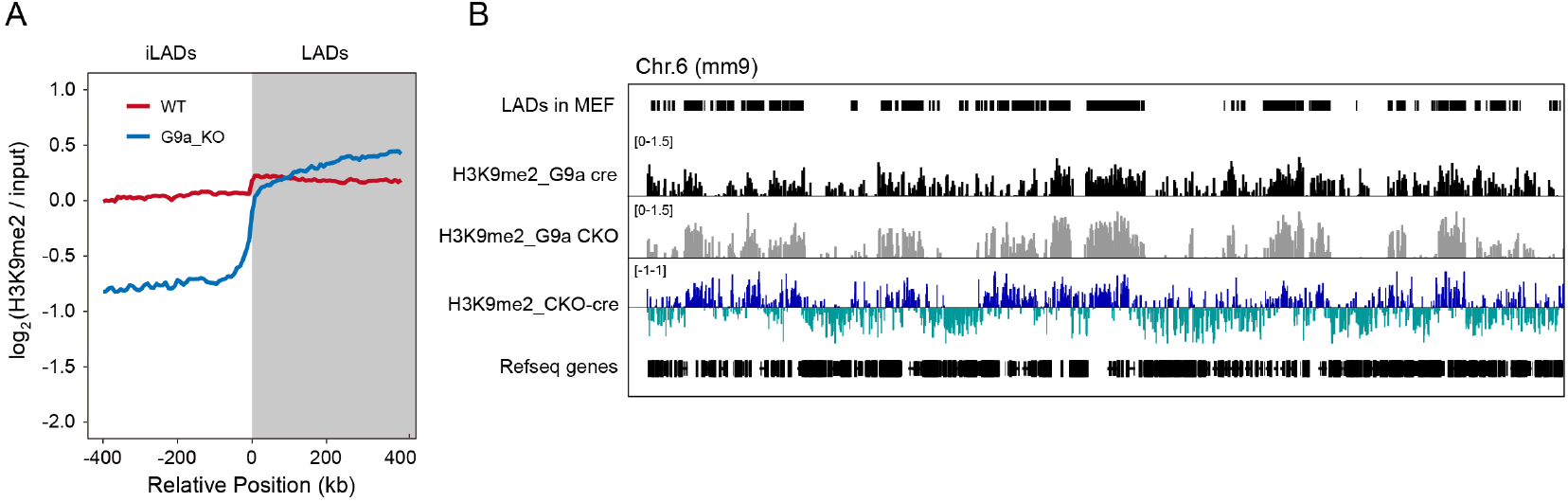
Region-specific removal of H3K9me2 in G9a-KO MEF cells and G9a-CKO cardiomyocytes of mouse. **A.** Aligned H3K9me2 profiles of mirrored border regions of LADs in WT and G9a knockout MEF cells. H3K9me2 ChIP-seq data were from Chen *et al.* [27]. **B.** Representative tracks of normalized H3K9me2 ChIP-seq in Cre or G9a-CKO cardiomyocytes; data from Papait *et al.* [28].

**Figure S3.**
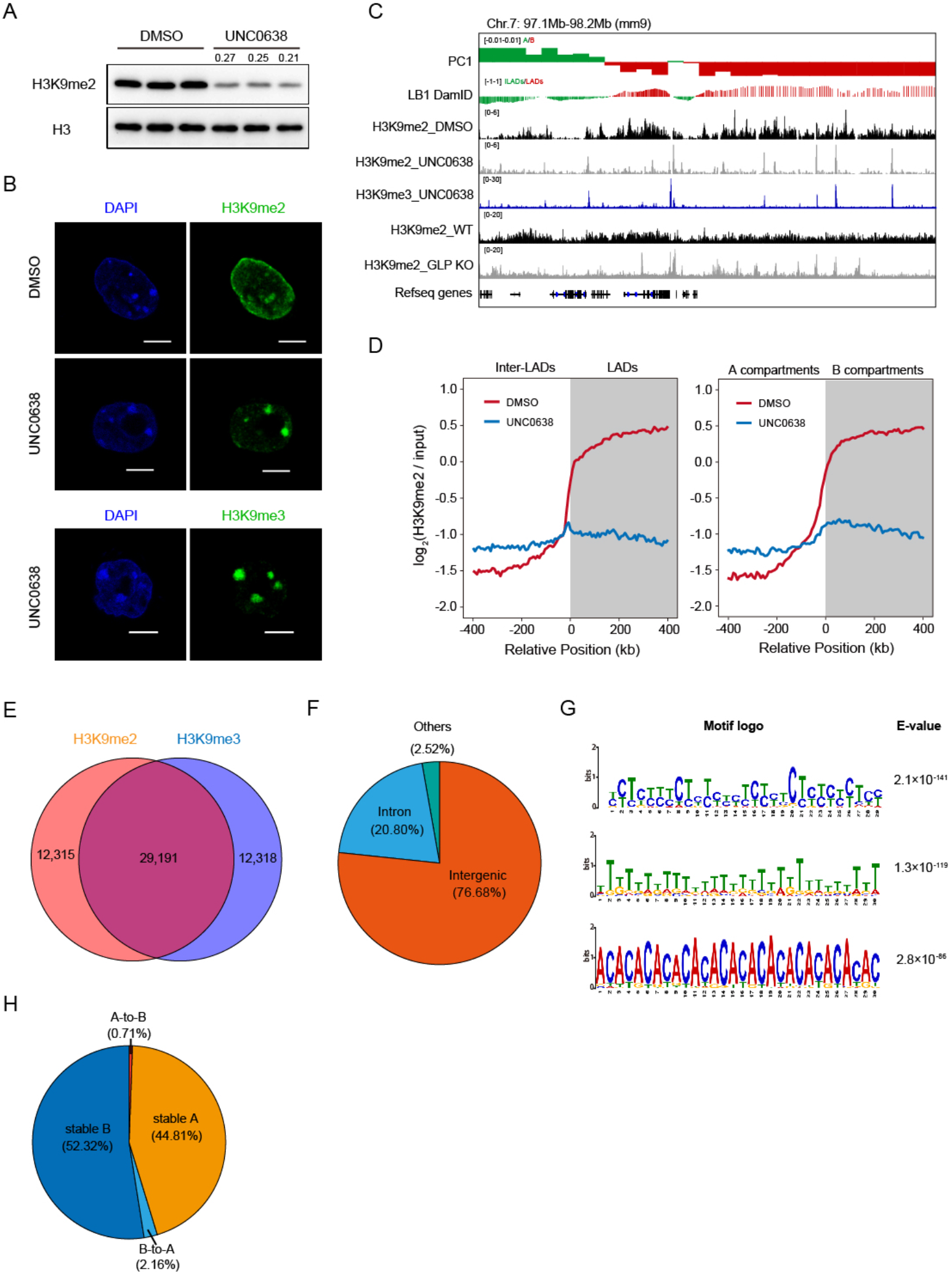
Inhibition of G9a/GLP removes most of H3K9me2 modifications in mESCs. **A.** Western blotting of H3K9me2 in DMSO and UNC0638 treated mESCs, with three biological repeats respectively (independent DMSO/UNC0638 treatment). H3 serves as the loading control. The numbers are relative grayscale of western blotting calculated by AlphaView software. **B.** Representative IF images of H3K9me2 in DMSO and UNC0638 treated mESC; H3K9me3 in UNC0638 treated mESCs. Scale bar, 5 μm. **C.** Representative H3K9me2 ChIP-seq tracks of DMSO/UNC0638 treated and WT/GLP KO mESCs; H3K9me3 ChIP-seq tracks of UNC0638 treated mESCs; PC1 values and smoothed Lamin B1 DamID signals of mESC. **D.** Aligned H3K9me2 profiles of mirrored border regions of LADs (left) and A/B compartments (right) in DMSO and UNC0638 treated mESCs. **E.** Venn diagram showing peaks overlapping between H3K9me2 and H3K9me3 ChIP-seq of UNC0638 treated mESCs. **F.** Proportions of the remaining H3K9me2 peaks not overlapped with H3K9me3 peaks in gene location of UNC0638 treated mESC. G. Motif analysis of the remaining H3K9me2 peaks not overlapped with H3K9me3 peaks in UNC0638 treated mESC. **H.** Compartment switching after UNC0638 treatment in mESCs.

**Figure S4.**
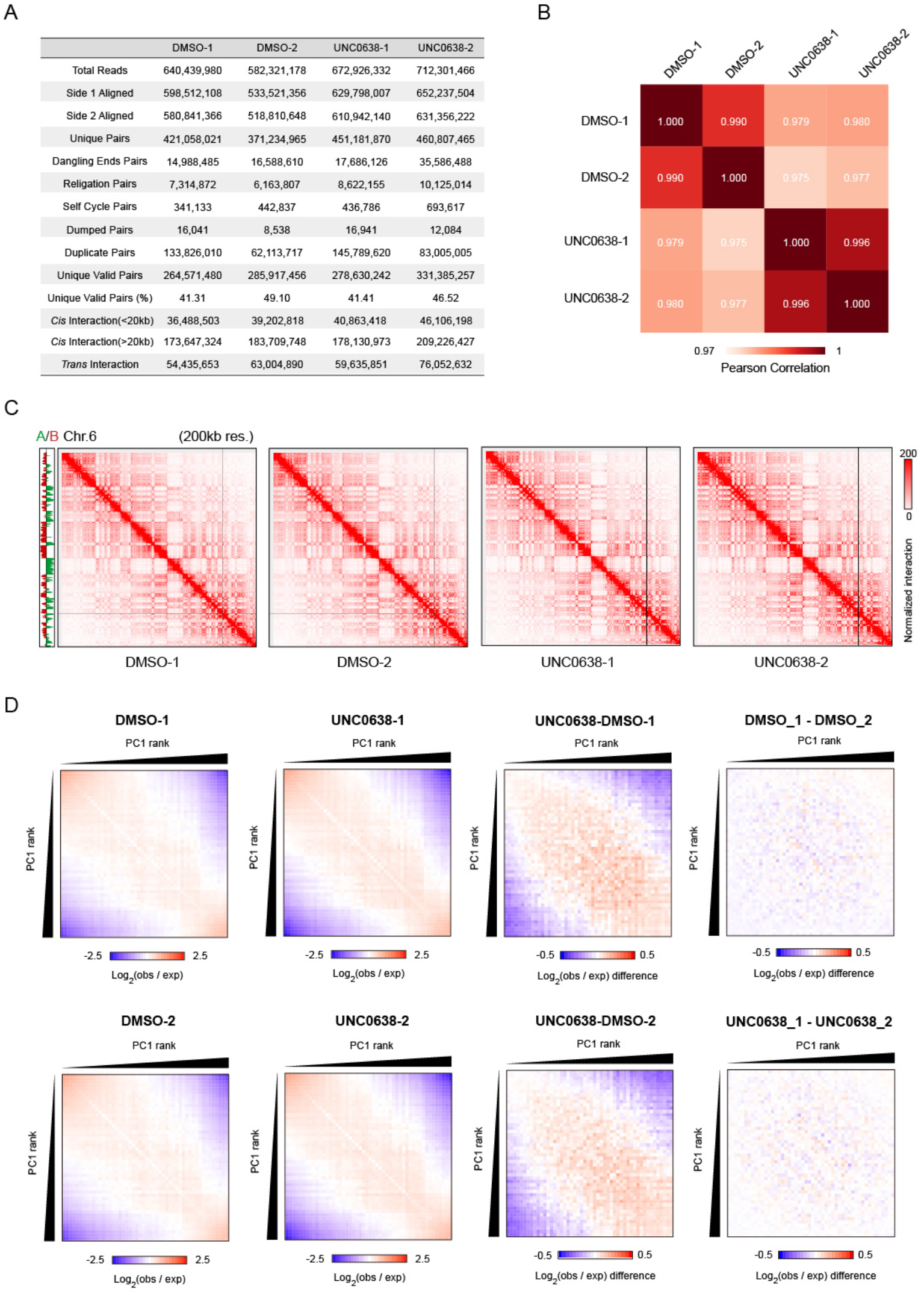
Mapping statistics and reproducibility of Hi-C experiments. **A.** Mapping statistics of Hi-C deep sequencing data of DMSO and UNC0638 treated AML12 cells, with two biological repeats respectively (independent treatments, Hi-C assay and sequencing). **B.** Pearson correlation coefficients of PC1 values derived from compartment analysis between replicates. **C.** Representative contact matrices of DMSO and UNC0638 treated AML12 cells. **D.** Average contact enrichment between pairs of 250 kb bins ranked by PC1 values in DMSO (1^st^ column) and UNC0638 (2^nd^ column) treated AML12 cells, and the difference between them (3^rd^ and 4^th^ columns), with two biological repeats respectively.

## Supplemental Tables (Separate files)

**Table S1** Coordinates of GSRs in AML12 cell (mm9).

**Table S2** LADs identified in DMSO and UNC0638 treated AML12 cells (mm9).

**Table S3** TAD boundaries identified in DMSO and UNC0638 treated AML12 cells (mm9).

**Table S4** Differentially expressed genes between DMSO and UNC0638 treated AML12 cells (mm10).

**Table S5** Genes identified in clusters 1-4 in AML12 cells (mm10).

